# Phenotypic associations among cell cycle genes in *Saccharomyces cerevisiae*

**DOI:** 10.1101/2020.02.13.948257

**Authors:** María Bermudez-Cruz, Peter I. Wu, Deanna Callerame, Staci Hammer, James C. Hu, Michael Polymenis

## Abstract

A long-standing effort in biology is to precisely define and group phenotypes that characterize a biological process, and the genes that underpin them. In *Saccharomyces cerevisiae* and other organisms, functional screens have generated rich lists of phenotypes associated with individual genes. However, it is often challenging to identify sets of phenotypes and genes that are most closely associated with a given biological process. Here, we focused on the 166 phenotypes arising from loss-of-function and the 86 phenotypes from gain-of-function mutations in 571 genes currently assigned to cell cycle-related ontologies in *S. cerevisiae*. To reduce this complexity, we applied unbiased, computational approaches of correspondence analysis to identify a minimum set of phenotypic variables that accounts for as much of the variability in the data as possible. Loss-of-function phenotypes can be reduced to 20 dimensions, while gain-of-function ones to 14 dimensions. We also pinpoint the contributions of phenotypes and genes in each set. The approach we describe not only simplifies the categorization of phenotypes associated with cell cycle progression but can also serve as a discovery tool for gene function.

## INTRODUCTION

The generation of systematic mutant collections in a variety of model systems enables large-scale phenotypic screens, which are now standard in academic and commercial settings. The first organism for which such mutant collections became available is the budding yeast *Saccharomyces cerevisiae* (Giaever and Nislow 2014). As a result, there is a wealth of phenotypes associated with most genes in that organism, displayed in easily accessible databases (Engel *et al.* 2009; Cherry *et al.* 2012). Gene Ontology (GO) techniques accurately specify the semantic relationships between terms, and they are indispensable for representing and organizing the accumulating biological knowledge (Ashburner *et al.* 2000). Curations of the literature and computational approaches have given rise to the systematic categorization of individual genes to biological processes.

However, given the numerous phenotypes often associated even with a single gene, the more genes involved in a biological process, the larger the number of phenotypes associated with that process. Hence, despite the plethora of phenotypic information on a per-gene basis, there is a loss in clarity and priority to the phenotypes most pertinent to the biological process in question. For example, at the time of preparing this report, based on the information on the Saccharomyces Genome Database (Cherry *et al.* 2012), there were at least 571 *S. cerevisiae* genes assigned to cell cycle related processes (see next Section). Collectively, there were 166 loss-of-function phenotypes associated with these genes, with additional qualifiers raising that number to 371 phenotypes. Among this bewildering set, identifying the phenotypic variables that cluster together in different groups and the genes that drive this classification may offer new insights into phenotype-phenotype and gene-phenotype associations within this biological process.

Network-based approaches have been used to link diseases with disease genes in humans, revealing common genetic origins of several conditions (Goh *et al.* 2007). Widely used multivariate statistical techniques can simplify related variables. Measuring the degree that the observed variables correlate with each other, provides the basis for the number of variables in a dataset to be reduced. If two or more phenotypic variables share some features, then based on the magnitude and direction of the relationship, the observed complexity may be simplified. Techniques implementing the above principles include factor analysis and principal component analysis (Child 1990). For categorical data (e.g., the presence or absence of a phenotype), a related approach is that of correspondence analysis (BenzeĆri 1992).

Here, we identified 571 genes associated with cell division and cell cycle progression. We applied correspondence analysis to examine the numerous phenotypes associated with these genes, resulting both from loss- and gain-of-function mutations. Some phenotypic associations were generic, with mutations affecting vegetative and respiratory growth, or resistance to toxins, pH, and metals. In other cases, the clustering of some phenotypes and the gene associations was consistent with the literature. For example, loss-of-function mutations that affect shmoo formation and mating efficiency together contributed most significantly in one of the dimensions. Likewise, gain-of-function mutations affecting cellular morphology, size, and budding index together contributed significantly in another dimension. Hence, systematic phenotypic associations provide a useful dissection of biological processes and gene functions.

## RESULTS

### Gene set

Before analyzing any phenotypes associated with cell division and cell cycle progression, it is essential to identify the genes related to these processes. At the time of writing this report, the biological process ‘cell cycle’ (GO:0007049) was defined as: “The progression of biochemical and morphological phases and events that occur in a cell during successive cell replication or nuclear replication events. Canonically, the cell cycle comprises the replication and segregation of genetic material followed by the division of the cell …” (https://www.yeastgenome.org/go/7049). There were 307 genes annotated to the ‘cell cycle’ biological process (File1). However, we noticed that some genes that govern vital cell cycle events were not in this set. For example, *SIC1*, encoding a cyclin-dependent kinase inhibitor that must be destroyed for DNA replication to begin. Destruction of Sic1p is the only essential function of G1 cyclins (Schneider *et al.* 1996). Another gene that was not in the computationally annotated ‘cell cycle’ genes was *MPS1*, which encodes a conserved kinase that is essential for spindle pole body duplication (Liu and Winey 2012).

Consequently, we included additional biological processes (File1), such as ‘DNA replication’ (GO:0006260), ‘chromosome segregation’ (GO:0007059), ‘cell division’ (GO:0051301). All the genes in the ‘cell division’ process were annotated computationally and were also in the ‘cell cycle’ set (Figure 1). However, several genes in the ‘DNA replication’ and ‘chromosome segregation’ processes, were not annotated as ‘cell cycle’ genes (Figure 1). We also noted that there was incomplete overlap between the genes that were annotated computationally or by manual curation within the ‘DNA replication’ and ‘chromosome segregation’ processes themselves (File1, sheets 0006260 and 0007059). To ensure that our list of cell cycle genes is as comprehensive as possible, we included all ‘children’ categories to the above gene ontology nodes. These additional categories (n=100) are listed in File1/sheet ‘categories’ (see also the individual sheets numbered as the corresponding gene ontologies), and they were grouped as ‘OTHER’ (see File1/sheet ‘sets_Fig1’). The overlap between the ‘cell cycle’ (GO:0007049), ‘DNA replication’ (GO:0006260), ‘chromosome segregation’ (GO:0007059), ‘cell division’ (GO:0051301), and ‘OTHER’ sets is shown in Figure 1. A total of 185 genes were unique to the ‘OTHER’ set. Overall, there were 571 unique genes in all these, gene ontology-based, biological processes related to cell division, and cell cycle progression (File1/sheet: ‘genes’). In the rest of this study, we analyzed the loss- and gain-of-function phenotypes associated with each of these 571 genes.

**FIGURE 1.**
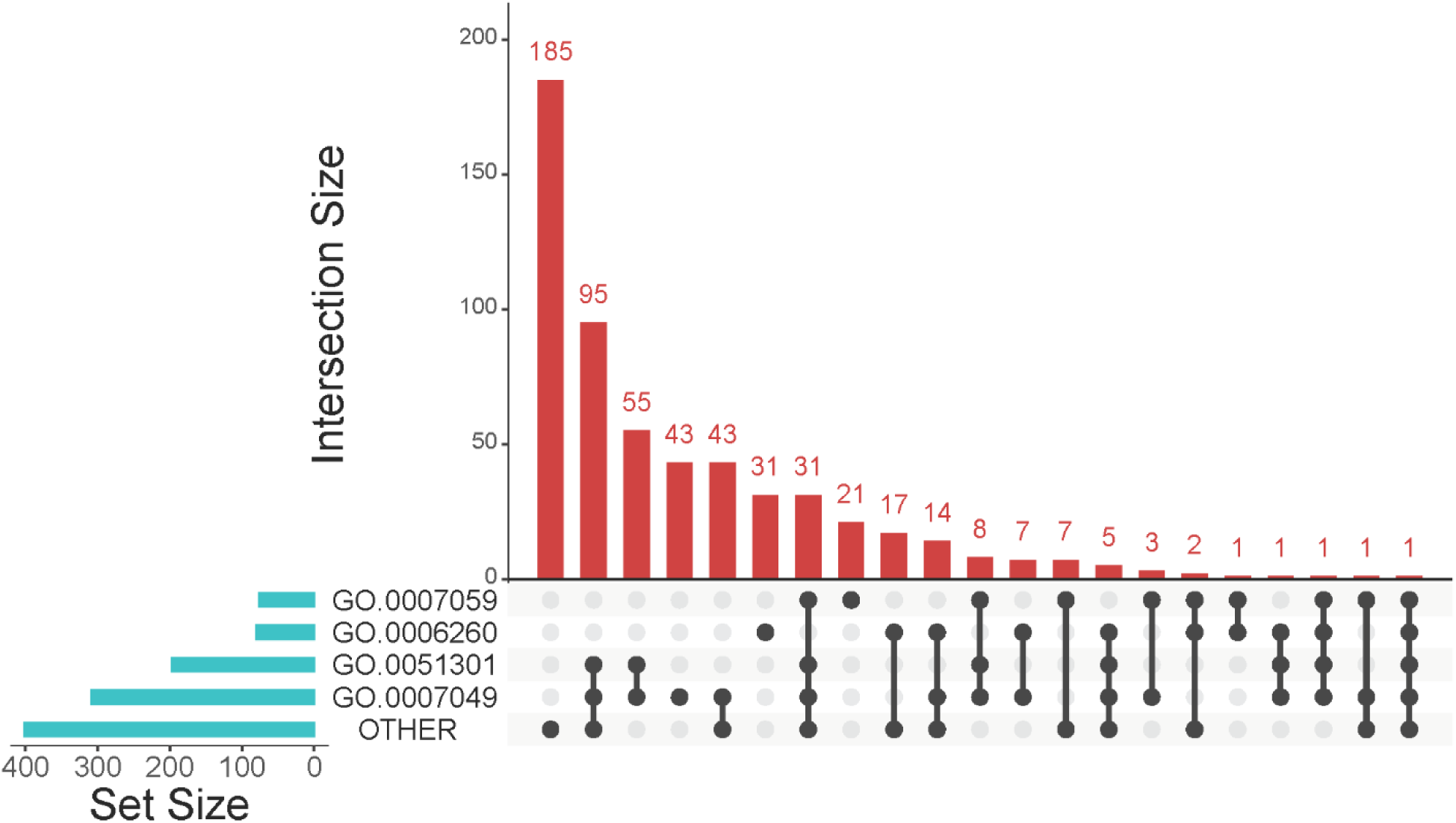
Gene ontologies related to cell cycle progression and cell division. Matrix layout for all intersections of the sets of genes we interrogated. The biological processes were ‘cell cycle’ (GO:0007049), ‘DNA replication’ (GO:0006260), ‘chromosome segregation’ (GO:0007059), ‘cell division’ (GO:0051301). In ‘OTHER’ there were genes grouped together from various cell cycle-related ontologies, as described in the text and in Materials and Methods. The size of the sets is shown on the bar plot to the left. The number of genes unique to the indicated intersections is shown separately on the bar plot to the right. The names of all genes in each set are shown in File1/sheet ‘sets_Fig1’. The graph was drawn with the *UpSet* R language package.

### Loss-of-function phenotypes

To analyze the 166 phenotypes associated with loss-of-function mutations in 561 genes, we tabulated them as we describe in the Materials and Methods. Correspondence analysis was performed with the R language package *FactoMiner*, and the related ones *factoextra* and *FactoInvestigate* (see Materials and Methods). We found that there were 20 significant dimensions, accounting for ≈2/3 of the observed variance (Figure 2, bottom). The percentage of the 561 genes associated with each of these 20 dimensions is shown in Figure 2, top. A detailed list is in File2/sheet ‘lof_gene_20dim’.

**FIGURE 2.**
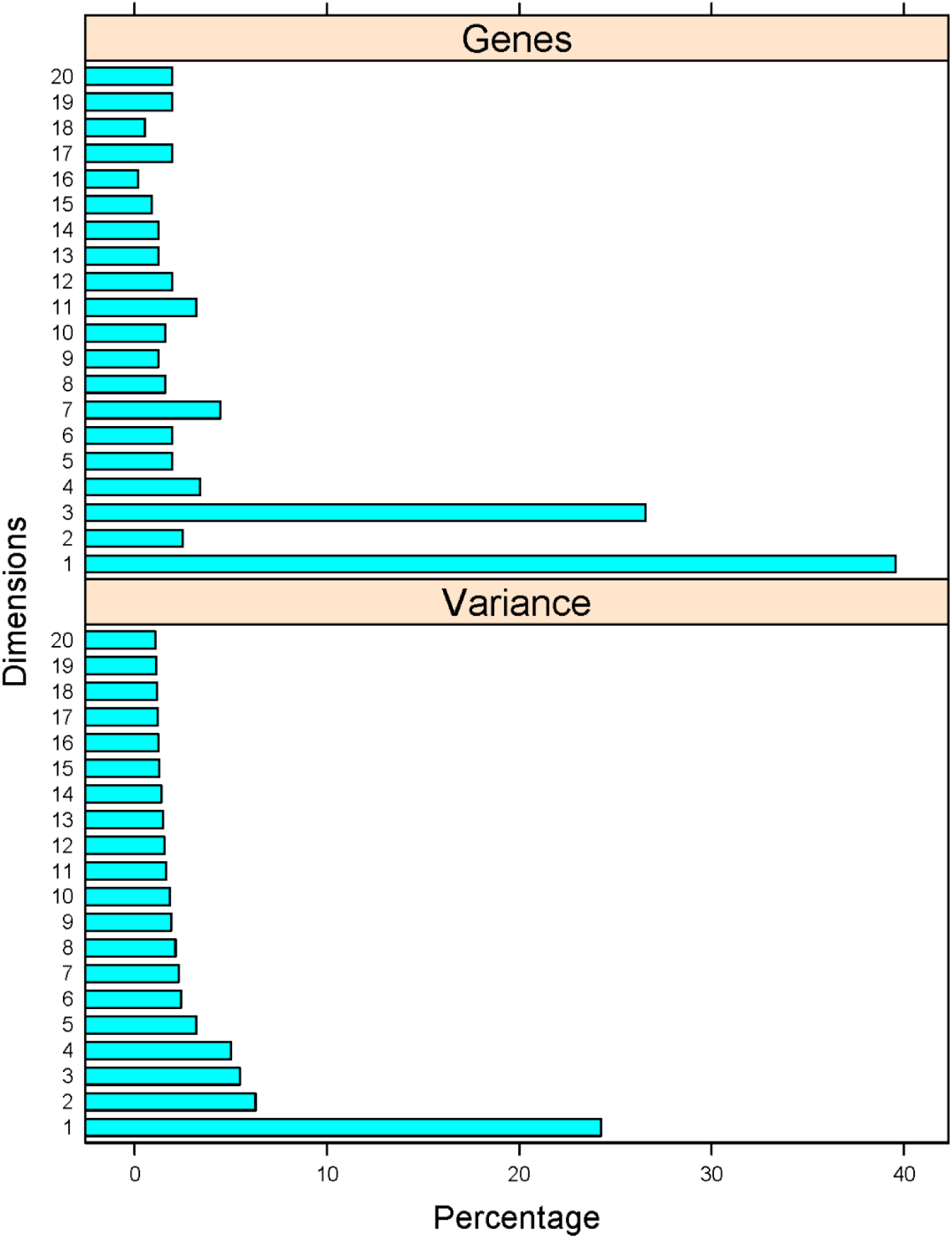
Phenotypic variance and gene associations with the 20 dimensions from the multiple correspondence analysis of the loss-of-function phenotypes of cell cycle-related genes. *Top*, The percentage of genes (x-axis) most closely associated with each of the dimensions (y-axis). *Bottom*, The percentage of the variance (x-axis) explained by each of the dimensions shown (y-axis).

A major objective is to identify which phenotypic variables the 20 dimensions are the most linked to, in other words which phenotypes describe the best each dimension. For the loss-of-function phenotypes, this is shown graphically in Figure 3 (detailed lists for each phenotype and dimension are in File2). The phenotypes that were most significantly associated (R^2^>0.2) with the most populous dimension (#1; 39% of all genes), were very general, and not particularly informative (Figure 3): chemical compound accumulation, respiratory or vegetative growth, metal resistance, etc (see File2/sheet ‘res1_dimdesc’). The only other cell cycle-related phenotype in this group was ‘cell size’. Cell size changes are often interpreted as perturbations in the normal coupling of cell growth with cell division (Jorgensen *et al.* 2002), albeit there is not a strong correlation between cell size and the length of the G1 phase of the cell cycle (Hoose *et al.* 2012; Blank *et al.* 2018). In other dimensions, interesting and expected associations were evident. For example, in Dimension 2, ‘shmoo formation’, ‘bud neck morphology’, and ‘pheromone induced cell cycle arrest’ were clustered together (Figure 3). Secretory processes were featured heavily in Dimension 4, with the phenotypes affecting ‘endoplasmic reticulum distribution’, ‘peroxisomal morphology’, ‘Golgi distribution’. Similarly, ‘vesicle distribution’ and ‘vacuolar transport’ were associated with Dimension 15. The constellation of phenotypes associated with loss-of-function mutations in *TOR2* is unique. *TOR2* is the only gene in Dimension 16, with ‘metabolism and growth’ and ‘osmotic stress resistance’ being the most prominent phenotypes. The remaining dimensions were defined by phenotypes that were only weakly (R^2^<0.2), or not directly, associated with cell cycle progression.

**FIGURE 3.**
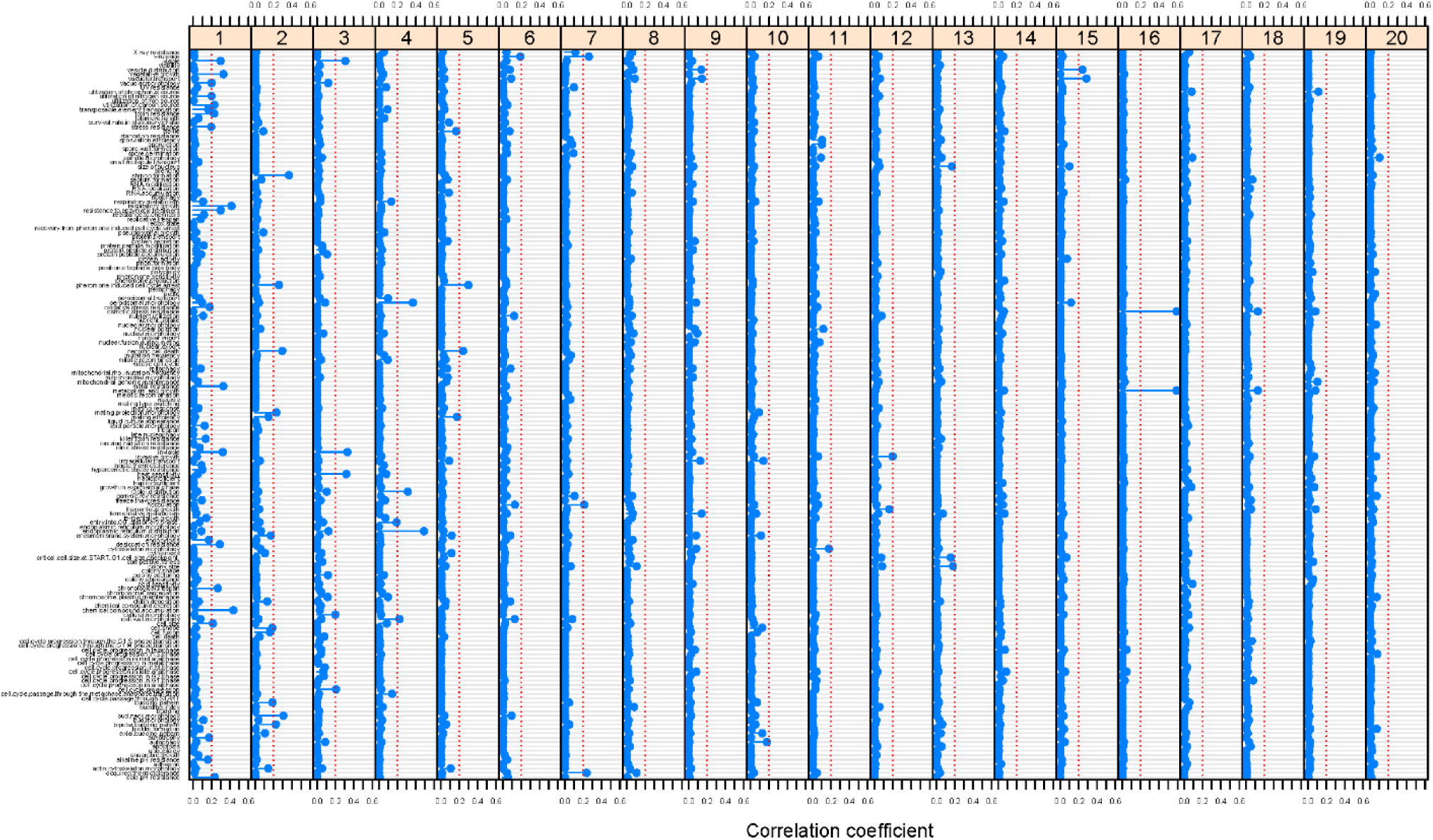
Correlation of the loss-of-function phenotypes with each of the 20 dimensions. Each panel corresponds to a dimension, and the associated correlation coefficients are shown on the x-axis. The dotted red line in each panel indicates the 0.2 R^2^ cutoff value.

### Gain-of-function phenotypes

There were 86 phenotypes associated with gain-of-function mutations in 368 genes (from a total of 571 genes). The phenotypic matrix was organized and analyzed as for the gain-of-function mutations (see Materials and Methods). Based on correspondence analysis we found that there were 14 significant dimensions (Figure 4, bottom), with the vast majority of genes grouped in just one dimension (#2; see Figure 4, top). A detailed list is in File3/sheet ‘gof_gene_14dim’.

**FIGURE 4.**
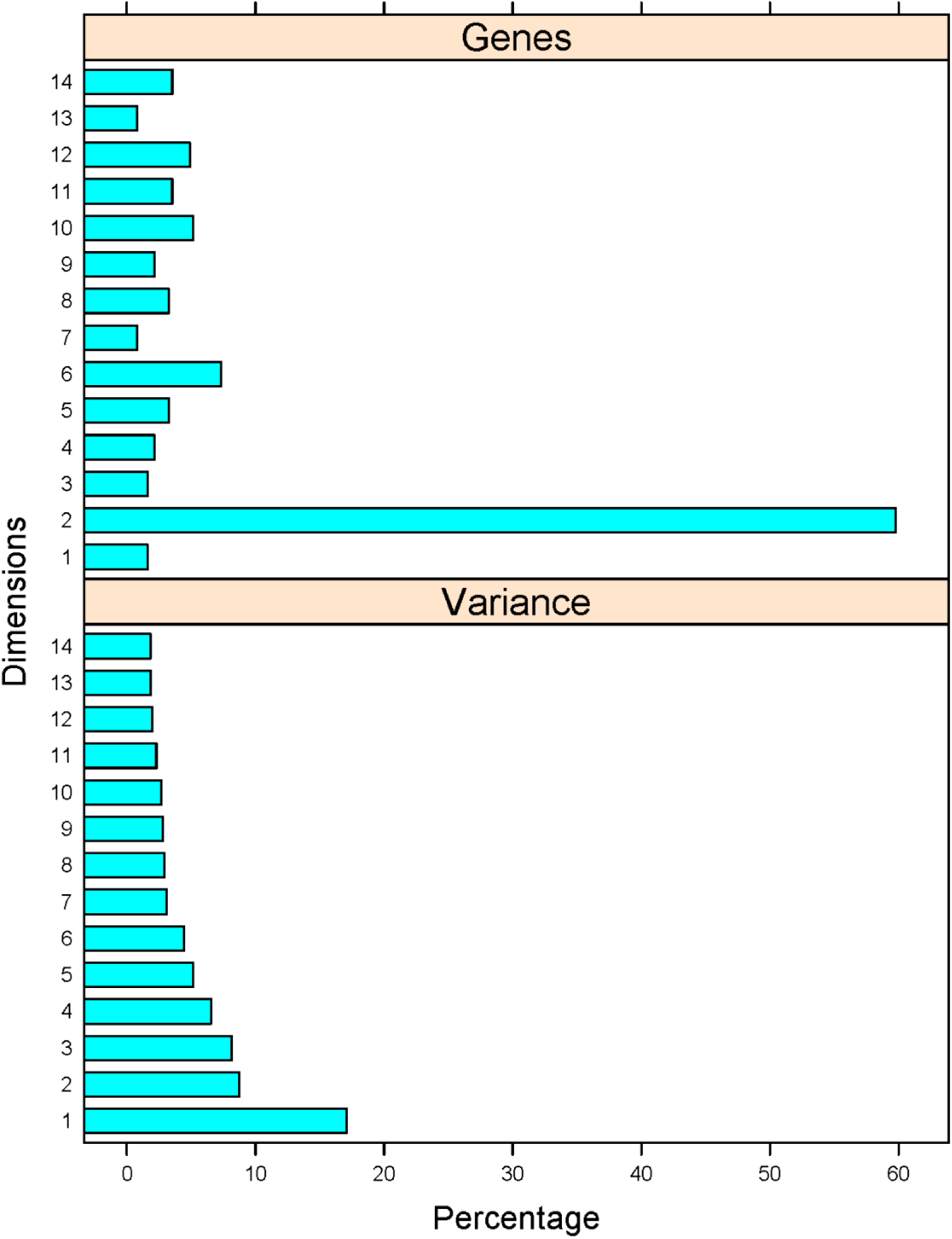
Phenotypic variance and gene associations with the 14 dimensions from the multiple correspondence analysis of the gain-of-function phenotypes of cell cycle-related genes. *Top*, The percentage of genes (x-axis) most closely associated with each of the dimensions (y-axis). *Bottom*, The percentage of the variance (x-axis) explained by each of the dimensions shown (y-axis).

We next identified the phenotypic variables for the gain-of-function mutants describe the best each dimension (Figure 5, detailed lists for each phenotype and dimension are in File3). Most genes (≈60%) were grouped in Dimension 2. The phenotypes that contributed most significantly (R^2^>0.2) to Dimension 2 were: ‘cellular morphology’, ‘budding index’ (a proxy for altered cell cycle progression), ‘cell size’, and ‘cell cycle progression in G2 phase’ (Figure 5).

**FIGURE 5.**
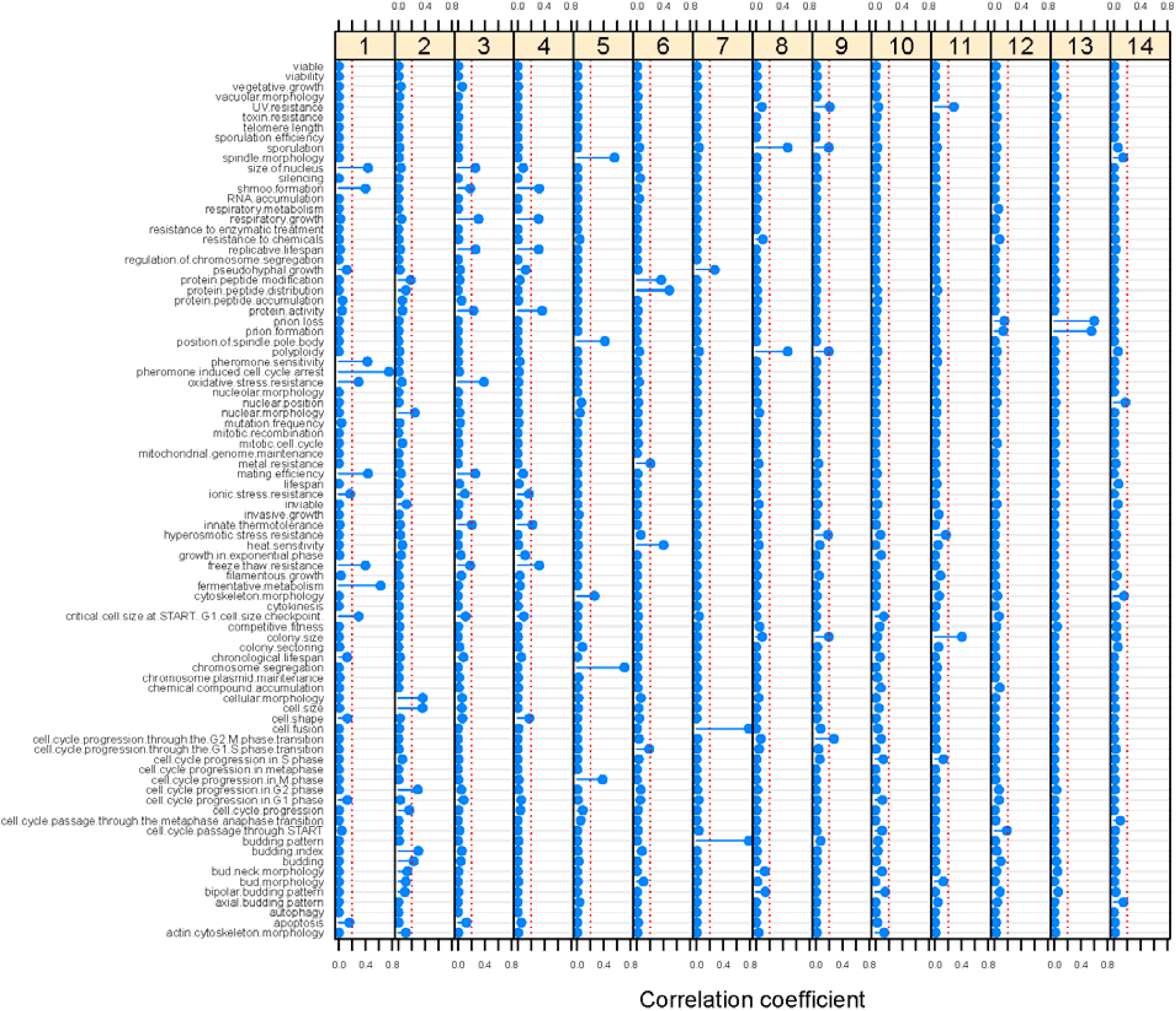
Correlation of the gain-of-function phenotypes with each of the 14 dimensions. Each panel corresponds to a dimension, and the associated correlation coefficients are shown on the x-axis. The dotted red line in each panel indicates the 0.2 R^2^ cutoff value.

The clustering of relevant phenotypes was also evident in other dimensions. For example, ‘chromosome segregation’, ‘spindle morphology’, ‘position of spindle pole body’, and ‘cell cycle progression in M phase’ were all strongly associated with Dimension 5. On the other hand, ‘pheromone induced cell cycle arrest’, ‘mating efficiency’, ‘pheromone sensitivity’, ‘shmoo formation’ were all clustered together in Dimension 1. In the same Dimension, we also noticed the phenotypes ‘size of nucleus’ and ‘critical cell size at START – G1 cell size checkpoint’. These are phenotypes associated with over-expression of the G1 cyclin Cln3p. The *CLN3* gene is most closely associated with Dimension 1 (see File 3/sheet ‘gof_gene_14dim’). We note that *CLN3* was originally identified not only on the basis of reduced cell size when over-expressed (Sudbery *et al.* 1980; Nash *et al.* 1988), but also because it can bypass the pheromone-induced cell cycle arrest (Cross 1988).

## DISCUSSION

The results we presented are significant for two reasons: First, the multitude of phenotypes associated with genes involved in cell cycle progression can be grouped in a smaller number of categories, simplifying their analysis and the gene contributions to each category. Second, the approach we described ought to apply to any biological process.

When testing gene function, the old maxim “when in doubt knock-it out” took a more expansive turn with the availability of genome-wide deletion sets. For several model systems, and especially *S. cerevisiae*, these sets enable large-scale, often automated, phenotypic assays (Giaever *et al.* 2002; Giaever and Nislow 2014). As the phenotypes associated with each gene increase, it becomes less clear which of the phenotypes associated with each gene are the most pertinent to the biological process in question. A key component in addressing this issue is high-quality annotation from the available databases. Gene Ontology (GO) categories standardize gene product annotations with regards to molecular function, biological process, and cellular component. *S. cerevisiae* is probably better annotated than most other experimental organisms, with computational and human-based approaches (Cherry *et al.* 2012). Yet, even in this organism, as we showed for the cell cycle genes (Figure 1), there is not a complete overlap among the different approaches, underscoring the need for continued efforts to improve systematic annotation (Siegele *et al.* 2019). Nonetheless, the existing information and curation efforts are invaluable, and formed the basis of our analysis. The relative simple approaches we used here to cluster the diverse phenotypes reported in the literature are scalable to other biological processes and genomes.

## ACKNOWLEDGEMENTS

This work was supported by NIH grant R01GM123139 to M.P.

## AUTHOR CONTRIBUTIONS

MP conceptualized the project. MB-C, PIW, JCH, and MP designed experiments. MB-C, PIW, DC, and SH analyzed the relevant data. MP processed the data and wrote the first draft of the manuscript. All authors were involved in the editing of the manuscript.

## MATERIALS AND METHODS

### Datasets

All the individual phenotypic reports for each gene were downloaded from the Saccharomyces Genome Database (https://www.yeastgenome.org/). Loss-of-function phenotypes included not only those reported for ‘null’ alleles, but also ‘conditional’, ‘repressible’, and ‘reduction of function’ ones. Gain-of-function phenotypes included ‘activation’, and ‘overexpression’. Phenotypes that arose from ‘unspecified’ alleles were excluded from the analysis. To assemble the individual files into a single spreadsheet, we used R language packages. The files were read using the *readr* package. For example, for the loss-of-function files, the command was: lof_files = list.files(path = ‘…’, pattern = “*.txt”, full.names = TRUE). Then, the individual files were assembled into a list, with the command: lof_list = lapply(lof_files, read_tsv). The list components were combined into a dataframe with the following command from the *dplyr* package: lof_parent_child <-bind_rows(lof_list, .id = NULL). The resulting spreadsheet is in File2/sheet ‘lof_parent_child’. There were 371 loss-of-function phenotypes associated with 561 genes. However, in many cases, the phenotypic terms included qualifiers. For example, for the parent term ‘vegetative growth’ there were qualifiers, such as ‘increased’, ‘increased rate’, etc. To simplify the analysis, we removed these qualifiers and focused only on the 161 parent, loss-of-function phenotypic terms. To split the parent terms from their qualifiers, we used the following command from the *tidyr* package: lof_parent <-separate(data = lof_parent_child, col = phenotypes_lof, into = c(“parent_ontology”, “child_ontology”), sep = “:”, remove = TRUE, convert = FALSE, extra = “warn”, fill = “warn”). The resulting spreadsheet is in File2/sheet ‘lof_parent’. For the gain-of-function phenotypes, the analogous spreadsheets are in File3/sheet ‘gof_parent_child’ and ‘gof_parent’.

To gauge whether phenotypic profiles for genes in the loss-of-function dataset (lof_parent.txt) associate with functions, for each gene pair, we calculated the semantic similarity based on Gene Ontology annotations (Yu *et al.* 2010). For this analysis, the R language package *infotheo* was used to calculate the mutual information-based similarity metric for all pairs of genes. Then, the R language package *GOSemSim* was used to calculate the semantic similarity between gene pairs based on the GO annotations of either molecular function, biological process or cellular component (Yu *et al.* 2010). Significantly higher semantic similarity was indeed observed between genes that have more similar phenotypic profiles (Figure S1).

### Factor analysis

Multiple correspondence analysis (MCA) was performed with the R language package *FactoMiner*, and the related ones *factoextra*, and *FactoInvestigate*. For the loss-of-function phenotypes, we used the lof_parent spreadsheet as input (File2/sheet ‘lof_parent’), after it was transposed, so that the phenotypic variables were columns and the genes rows. Then we used the command: lof_MCA <-MCA(lof_parent, method = “Burt”). All the Eigen values associated with the analysis are in File2/sheet ‘lof_eigen’. To identify the number of the most significant dimensions, we used the command: dimRestrict(lof_MCA), which identified 20 dimensions as the most significant. We then re-run the MCA function for 20 dimensions, as follows: lof_MCA <-MCA(lof_parent, method = “Burt”, ncp = 20). The cosine values from the correspondence analysis represent the correlation coefficients (Child 1990). The cos2 values for the phenotypic variables were obtained with the command ‘get_mca_var(lof_MCA)’ and listed in File2/sheet ‘lof_var_cos2_20dim’. The cos2 values for the individuals (genes) were obtained with the command ‘get_mca_ind(lof_MCA)’ and they are listed in File2/sheet ‘lof_ind_cos2_20dim’. Based on this analysis, each of the genes was assigned to one of the 20 most significant dimensions (shown in File2/sheet ‘lof_gene_20dim’).

To interpret the dimensions, we used the ‘dimdesc’ function of the *FactoMiner* R language package. For each dimension (the example is for dimension 1), we run the command: res1_dimdesc = dimdesc(lof_MCA, axes=1:1, proba=1). The results for each dimension, with the R^2^ values for each phenotype and the associated p-value, are in the sheets of File2 (e.g., ‘res1_dimdesc’ for dimension 1, and so on). We grouped all the R^2^ values for each of the 166 phenotypes and 20 dimensions (File2/sheet ‘R2s’), used as input for Figure 3.

The analogous analysis was done for the gain-of-function phenotypes, and all the data are in File3.

## SUPPLEMENTARY FIGURES

**FIGURE S1.**
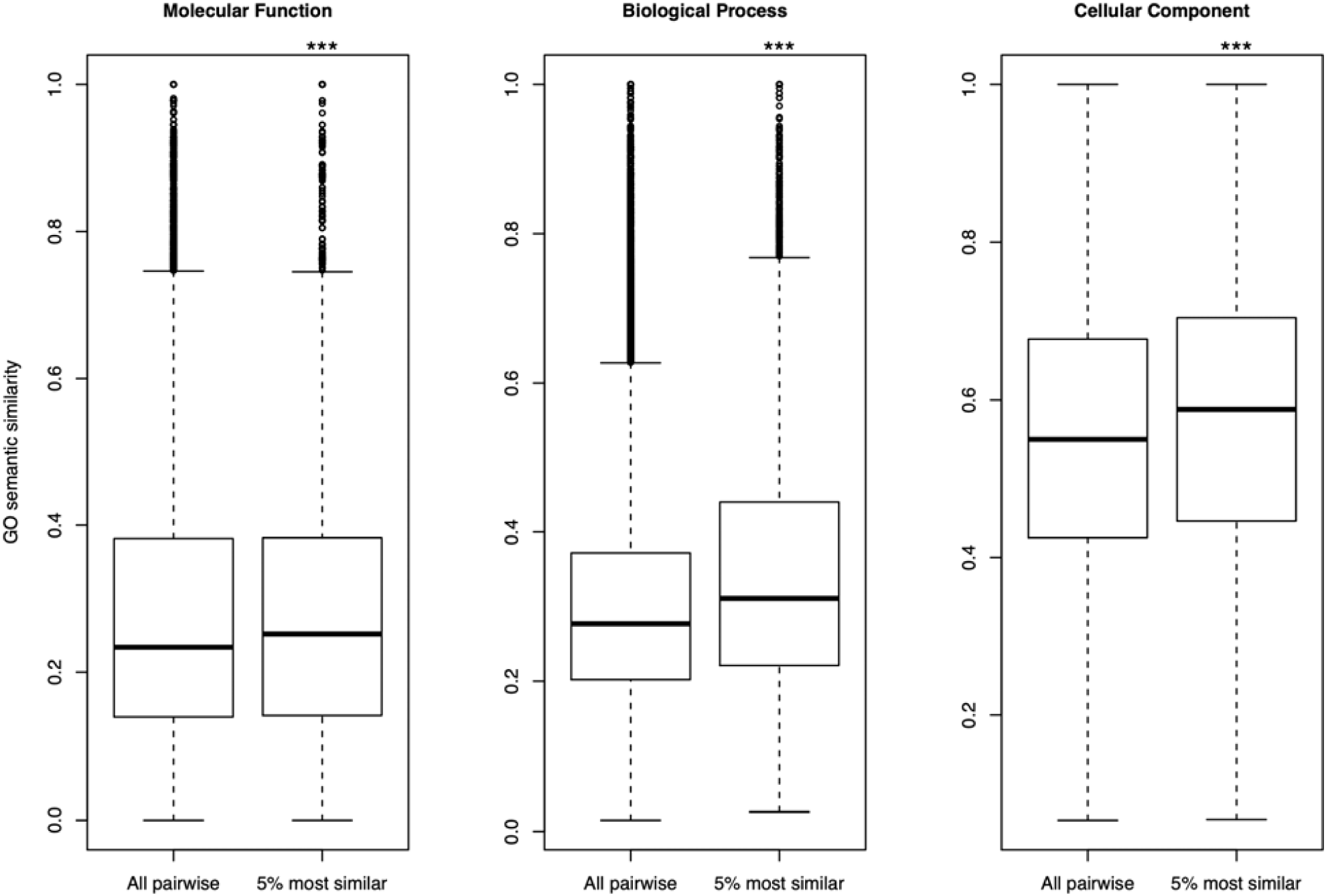
Higher semantic similarity between genes that have similar phenotypic profiles. Based on the SGD annotations that generated the phenotypic profiles, the top 5% phenotypically similar gene pairs have higher semantic similarity. ***: p-value<0.001 based on 1-sided Mann-Whitney U test.

